# An Effective Model of the Retinoic Acid Induced HL-60 Differentiation Program

**DOI:** 10.1101/138784

**Authors:** Ryan Tasseff, Holly A. Jensen, Johanna Congleton, Wei Dai, Katharine V. Rogers, Adithya Sagar, Rodica P. Bunaciu, Andrew Yen, Jeffrey D. Varner

## Abstract

In this study, we present an effective model All-Trans Retinoic Acid (ATRA)-induced differentiation of HL-60 cells. The model describes reinforcing feedback between an ATRA-inducible signalsome complex involving many proteins including Vav1, a guanine nucleotide exchange factor, and the activation of the mitogen activated protein kinase (MAPK) cascade. We decomposed the effective model into three modules; a signal initiation module that sensed and transformed an ATRA signal into program activation signals; a signal integration module that controlled the expression of upstream transcription factors; and a phenotype module which encoded the expression of functional differentiation markers from the ATRA-inducible transcription factors. We identified an ensemble of effective model parameters using measurements taken from ATRA-induced HL-60 cells. Using these parameters, model analysis predicted that MAPK activation was bistable as a function of ATRA exposure. Conformational experiments supported ATRA-induced bistability. Additionally, the model captured intermediate and phenotypic gene expression data. Knockout analysis suggested Gfi-1 and PPAR$[gamma] were critical to the ATRA-induced differentiation program. These findings, combined with other literature evidence, suggested that reinforcing feedback is central to hyperactive signaling in a diversity of cell fate programs.

## Introduction

Differentiation induction chemotherapy (DIC), using agents such as the vitamin A derivative all-trans retinoic acid (ATRA), is a promising approach for the treatment of many cancers^1–5^. For example, ATRA treatment induces remission in 80-90% of promyelocytic leukemia (APL) PML-RARα-positive patients^6^, thereby transforming a fatal diagnosis into a manageable disease^7^. However, remission is sometimes not durable and relapsed cases exhibit emergent ATRA resistance^8^, ^9^. To understand the basis of this resistance, we must first understand the ATRA-induced differentiation program. Toward this challenge, lessons learned in model systems, such as the lineage-uncommitted human myeloblastic cell line HL-60 reported to closely resemble patient derived cells^10^, could inform our analysis of the differentiation programs occurring in patients. Patient derived HL-60 leukemia cells have been a durable experimental model since the 1970’s to study differentiation^11^. HL-60 undergoes cell cycle arrest and either myeloid or monocytic differentiation following stimulation; ATRA induces G1/G0-arrest and myeloid differentiation in HL-60 cells, while 1,25-dihydroxy vitamin D3 (D3) induces arrest and monocytic differentiation. Commitment to cell cycle arrest and differentiation requires approximately 48 hr of treatment, during which HL-60 cells undergo two division cycles.

Sustained mitogen-activated protein kinase (MAPK) pathway activation is a defining feature of ATRA-induced HL-60 differentiation. ATRA drives sustained activation of the Raf/MEK/ERK pathway, leading to arrest and differentiation^12^. Betraying a feedback loop, MEK inhibition results in the loss of ERK as well as Raf phosphorylation and the failure to arrest and differentiate in response to ATRA^13^. Retinoic acid (and its metabolites) are ligands for the hormone activated nuclear transcription factors retinoic acid receptor (RAR) and retinoid X receptor (RXR)^14^. RAR/RXR activation is necessary for ATRA-induced Raf phosphorylation^13^ in concert with the formation of an ATRA-induced signalsome complex at the membrane, which drives MAPK activation. While the makeup of the signalsome complex is not yet known, we do know that it is composed of Src family kinases Fgr and Lyn, PI3K, c-Cbl, Slp76, and KSR, plus transcription factors AhR and IRF1^15–19^. Signalsome activity is driven by ATRA-induced expression of CD38 and putatively the heterotrimeric Gq protein-coupled receptor BLR1^20,21^. BLR1 (also known as CXCR5), identified as an early ATRA (or D3)-inducible gene using differential display^22^, is necessary for MAPK activation and differentiation^21^, and drives signalsome activity. Studies of the BLR1 promoter identified a non-canonical RARE site consisting of a 17 bp GT box approximately 1 kb upstream of the transcriptional start that conferred ATRA responsiveness^21^. Members of the BLR1 transcriptional activator complex, e.g. NFATc3 and CREB, are phosphorylated by ERK, JNK or p38 MAPK family members suggesting positive feedback between the signalsome and MAPK activation^23,24^. BLR1 overexpression enhanced Raf phosphorylation and accelerated terminal differentiation, while Raf inhibition reduced BLR1 expression and ATRA-induced terminal differentiation^25^. In particular, Raf phosphorylation of the NFATc3 transcription factors at the BLR1 promoter enables transcriptional activation at the RARE by ATRA bound to RAR/RXR^26^. BLR1 knock-out cells failed to activate Raf or differentiate in the presence of ATRA^25^. Interestingly, both the knockdown or inhibition of Raf, also reduced BLR1 expression and functional differentiation^25^. Thus, the expression of signalsome components e.g., BLR1 was Raf dependent, while Raf activation depended upon the signalsome. A previous computational study of ATRA-induced differentiation of HL-60 cells suggested that the BLR1-MAPK positive feedback circuit was sufficient to explain ATRA-induced sustained MAPK activation, and the expression of a limited number of functional differentiation markers^27^. Model analysis also suggested that Raf was the most distinct of the MAPK proteins. However, this previous study developed and analyzed a complex model, thus leaving open the critical question of what is the minimal positive feedback circuit required to drive ATRA-induced differentiation.

In this study, we explored this question using a minimal mathematical model of the key architectural feature of ATRA induced differentiation of HL-60 cells, namely positive feedback between an ATRA-inducible signalsome complex and MAPK activation. The ATRA responsive signalsome-MAPK circuit was then used to drive a down-stream gene expression program which encoded for the expression of intermediate and functional differentiation markers. The effective model used a novel framework which integrated logical rules with kinetic modeling to describe gene expression and protein regulation, while largely relying upon biophysical parameters from the literature. This formulation significantly reduced the size and complexity of the model compared to the previous study of Tasseff et al., while increasing the breadth of the biology described^27^. The effective model, despite its simplicity, captured key features of ATRA induced differentiation of HL-60 cells. Model analysis predicted the bistability of MAPK activation as a function of ATRA exposure; conformational experiments supported ATRA-induced bistability. Model simulations were also consistent with measurements of the influence of MAPK inhibitors, and the failure of BLR1 knockout cells to differentiate when exposed to ATRA. In addition, the expression of intermediate and phenotypic differentiation markers as also captured following ATRA exposure. Lastly, we showed through immunoprecipitation and inhibitor studies, that the guanine nucleotide exchange factor Vav1 is potentially a new ATRA-inducible member of the signalsome complex functioning as a regulator that contributes to signal amplification in the signalsome. Taken together, these findings when combined with other literature evidence, suggested that reinforcing feedback was central to differentiation programs generally, and necessary for ATRA-induced differentiation. The model answers a biologically important question that is not easily experimentally attacked, namely given the complexity of the signaling machine and the pathways it embodies, is there a critical small suite of molecules that are the action elements seminal to eliciting ATRA-induced cell differentiation and G0 arrest.

## Results

We constructed an effective model of ATRA-induced HL-60 differentiation which described signaling and gene expression events following the addition of ATRA (Fig. 1). The model connectivity was developed from literature and the studies presented here (Table 1). We decomposed the ATRA program into three modules; a signal initiation module that sensed and transformed the ATRA signal into activated cRaf-pS621 and the ATRA-RAR/RXR (Trigger) signals (Fig. 1A); a signal integration module that controlled the expression of upstream transcription factors given cRaf-pS621 and activated Trigger signals (Fig. 1B); and a phenotype module which encoded the expression of functional differentiation markers from the ATRA-inducible transcription factors (Fig. 1C). In particular, Trigger (a surrogate for the RARα/RXR transcriptional complex) regulated the expression of the transcription factors CCATT/enhancer binding protein α (C/EBPα), PU.1, and Egr-1. In turn, these transcription factors, in combination with cRaf-pS621, regulated the expression of downstream phenotypic markers such as CD38, CD11b or p47Phox. Each component of these modules was described by a mRNA and protein balance equation. Additionally, the signal initiation module also described the abundance of activated species e.g., Trigger and cRaf-pS621 whose values were derived from unactivated Trigger and cRaf protein levels. Lastly, because the population of HL-60 cells was dividing, we also considered a dilution term in all balance equations. The signal initiation module contained nine differential equations, while the signal integration and phenotype modules were collectively encoded by 54 differential equations. Model parameters were taken from literature (Table 2), or estimated from experimental data using heuristic optimization (see materials and methods).

**Figure 1.**
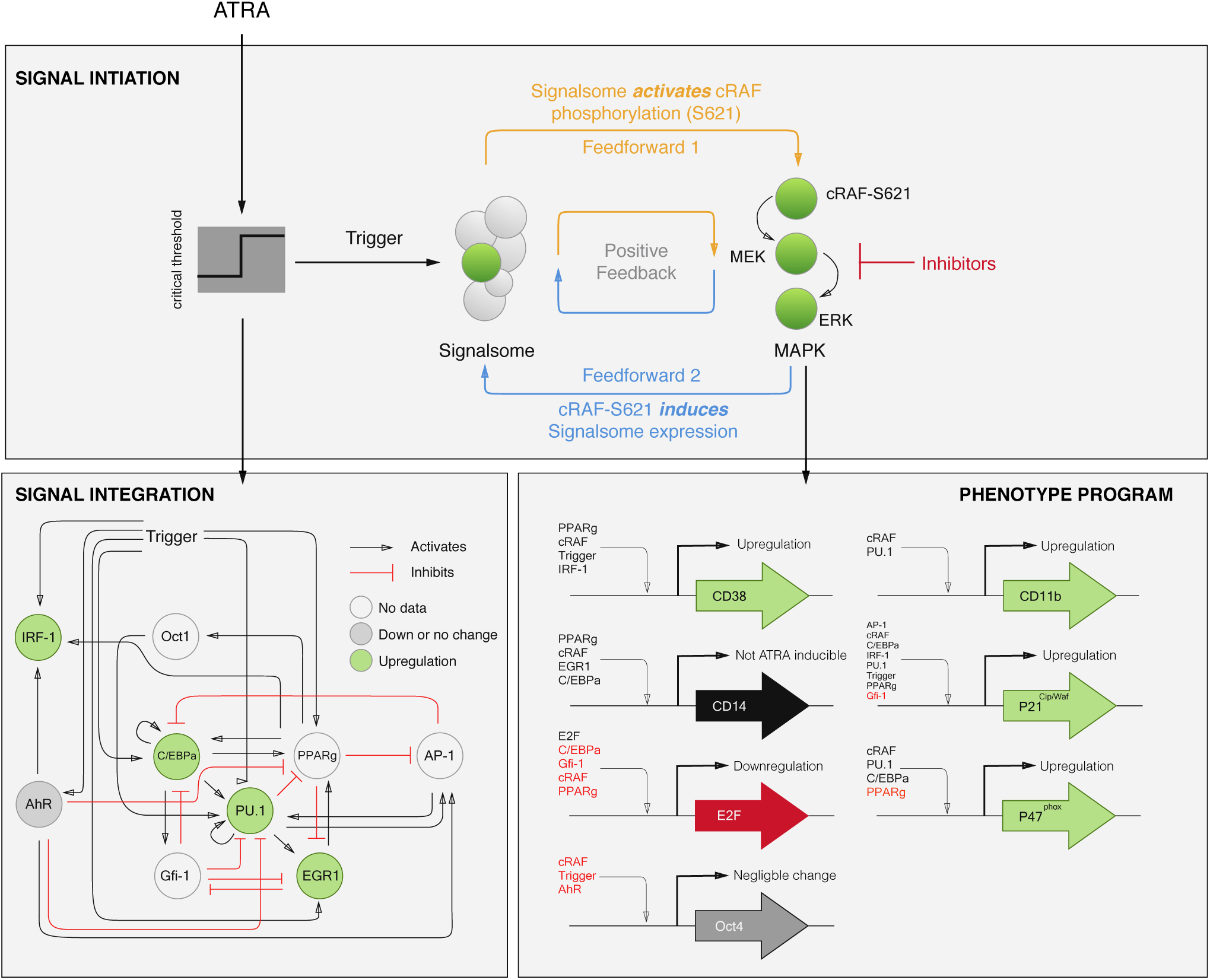
Schematic of the effective ATRA differentiation circuit. Above a critical threshold, ATRA activates an upstream Trigger, which induces signalsome complex formation. Signalsome activates the mitogen-activated protein kinase (MAPK) cascade which in turn drives the differentiation program and signalsome formation. Both Trigger and activated cRaf-pS621 drive a phenotype gene expression program responsible for differentiation. Trigger activates the expression of a series of transcription factors which in combination with cRaf-pS621 result in phenotypic change.

**Table 1.** Myelomonocytic transcription factor connectivity used in the signal integration and phenotype modules.

| Effector | Effect | Target | Source |
| --- | --- | --- | --- |
| RARα | + | RARα | 70 |
|  | + | PU.1 | 71 |
| | + | C/EBPα | 42 |
|  | + | IRF-1 | 72 |
|  | - | Oct4 | 73 |
|  | + | CD38 | 74 |
|  | + | p21 | 75 |
|  | + | AhR | 76 |
|  | + | Egr-1 | 77 |
| PPARγ | + | C/EBPα | 78 |
|  | + | IRF-1 | 79 |
|  | + | Oct1 | 80 |
|  | - | AP-1 | 81 |
|  | - | E2F | 82 |
|  | - | Egr-1 | 83 |
|  | + | CD38 | 84 |
|  | + | CD14 | 85 |
|  | + | p21 | 86 |
|  | - | p47Phox | 87 |
| PU.1 | - | PPARγ | 88 |
|  | + | PU.1 | 89 |
|  | + | AP-1 | 90 |
|  | + | Egr-1 | 38 |
|  | + | CD11b | 91 |
|  | + | p21 | 92 |
|  | + | p47Phox | 93 |
| C/EBPα | + | PPARγ | 78 |
|  | + | PU.1 | 43 |
| | + | C/EBPα | 94 |
|  | + | Gfi-1 | 95 |
|  | - | E2F | 96 |
|  | + | CD14 | 97 |
|  | + | p21 | 98 |
| IRF-1 | + | CD38 | 99 |
|  | + | p21 | 100 |
|  | - | PU.1 | 101 |
| | - | C/EBPα | 102 |
|  | - | E2F | 102 |
|  | - | Egr-1 | 38 |
|  | - | p21 | 102 |
| Oct1 | + | PU.1 | 103 |
| AP-1 | - | PPARγ | 81 |
|  | + | PU.1 | 104 |
|  | + | p21 | 105 |
| E2F | + | E2F | 106 |
| Egr-1 | + | PPARγ | 107 |
|  | - | Gfi-1 | 108 |
|  | + | CD14 | 109 |
| AhR | + | AP-1 | 110 |
|  | + | IRF-1 | 111 |
|  | - | Oct4 | 112 |
|  | - | PU.1 | 113 |

**Table 2.** Characteristic model parameters estimated from literature.

| Symbol | Description | Value | Units | Source |
| --- | --- | --- | --- | --- |
| $R_1$ | RNA polymerase abundance | 85,000 | copies/cell | 114, 115 |
| $R_2$ | Ribosome abundance | $1 \times 10^6$ | copies/cell | 116 |
| $G_i$ | Characteristic gene abundance | 2 | copies/cell | this study |
| $K_X$ | Saturation constant transcription | 600 | copies/cell | this study |
| $K_T$ | Saturation constant translation | 95,000 | copies/cell | this study |
| $t_{1/2,m}$ | characteristic mRNA half-life (transcription factor) | 2-4 | hr | 117 |
| $t_{1/2,p}$ | characteristic protein half-life | 10 | hr | 118 |
| $\theta_{m,j}$ | characteristic mRNA degradation constant | 0.34 | $\text{hr}^{-1}$ | derived |
| $\theta_{p,j}$ | characteristic protein degradation constant | 0.07 | $\text{hr}^{-1}$ | derived |
| $t_d$ | HL-60 doubling time | 19.5 | hr | this study |
| $\mu$ | growth rate | 0.035 | $\text{hr}^{-1}$ | derived |
| $k_d$ | death rate | $0.10\mu$ | $\text{hr}^{-1}$ | derived |
| $e_T$ | elongation rate RNA polymerase | 50-100 | nt/s | 119-122 |
| $e_X$ | elongation rate Ribosome | 5 | aa/s | 123 |
| $L_{T,o}$ | characteristic gene length | 44,192 | nt | 124 |
| $L_{X,o}$ | characteristic transcript length | 1,374 | nt | derived |
| $k_T$ | characteristic transcription rate | 1.44 | $\text{hr}^{-1}$ | derived |
| $k_X$ | characteristic translation rate | 3.60 | $\text{hr}^{-1}$ | derived |
| $k_{+,cRaf}$ | characteristic cRaf-S621 activation rate constant | 1 | $\text{hr}^{-1}$ | this study |
| $K_{+,cRaf}$ | characteristic saturation constant governing cRaf-pS621 formation | 60 | nM | this study |
| $K_D$ | characteristic MAPK inhibitor affinity | 10 | nM | this study |
| $D$ | Diameter of an HL-60 cell | 12.4 | $\mu\text{m}^3$ | 125 |
| $f_C$ | cytoplasmic fraction | 0.51 | dimensionless | 125 |

The signal initiation module recapitulated sustained signalsome and MAPK activation following exposure to 1*μ*M ATRA (Fig. 2A-B). An ensemble of effective model parameters was estimated by minimizing the difference between simulations and time-series measurements of BLR1 mRNA and cRaf-pS621 following the addition of 1*μ*M ATRA. We focused on the S621 phosphorylation site of cRaf since enhanced phosphorylation at this site is a defining characteristic of sustained MAPK signaling activation in HL-60. The effective model captured both ATRA-induced BLR1 expression (Fig. 2A) and sustained phosphorylation of cRaf-pS621 (Fig. 2B) in a growing population of HL-60 cells. Together, the reinforcing feedback within the signalsome and its embedded MAPK signaling axis led to sustained activation over multiple cellular generations. However, the effective model failed to capture the decline of BLR1 message after 48 hr of ATRA exposure. This suggested that we captured the logic leading to the onset of differentiation, but failed to describe program shutdown. Much of the focus in the literature has been on understanding the initiation of differentiation, with little attention paid to understanding how a program is terminated. This is a potential new direction that could be explored. Next, we tested the response of the signal initiation module to different ATRA dosages.

**Figure 2.**
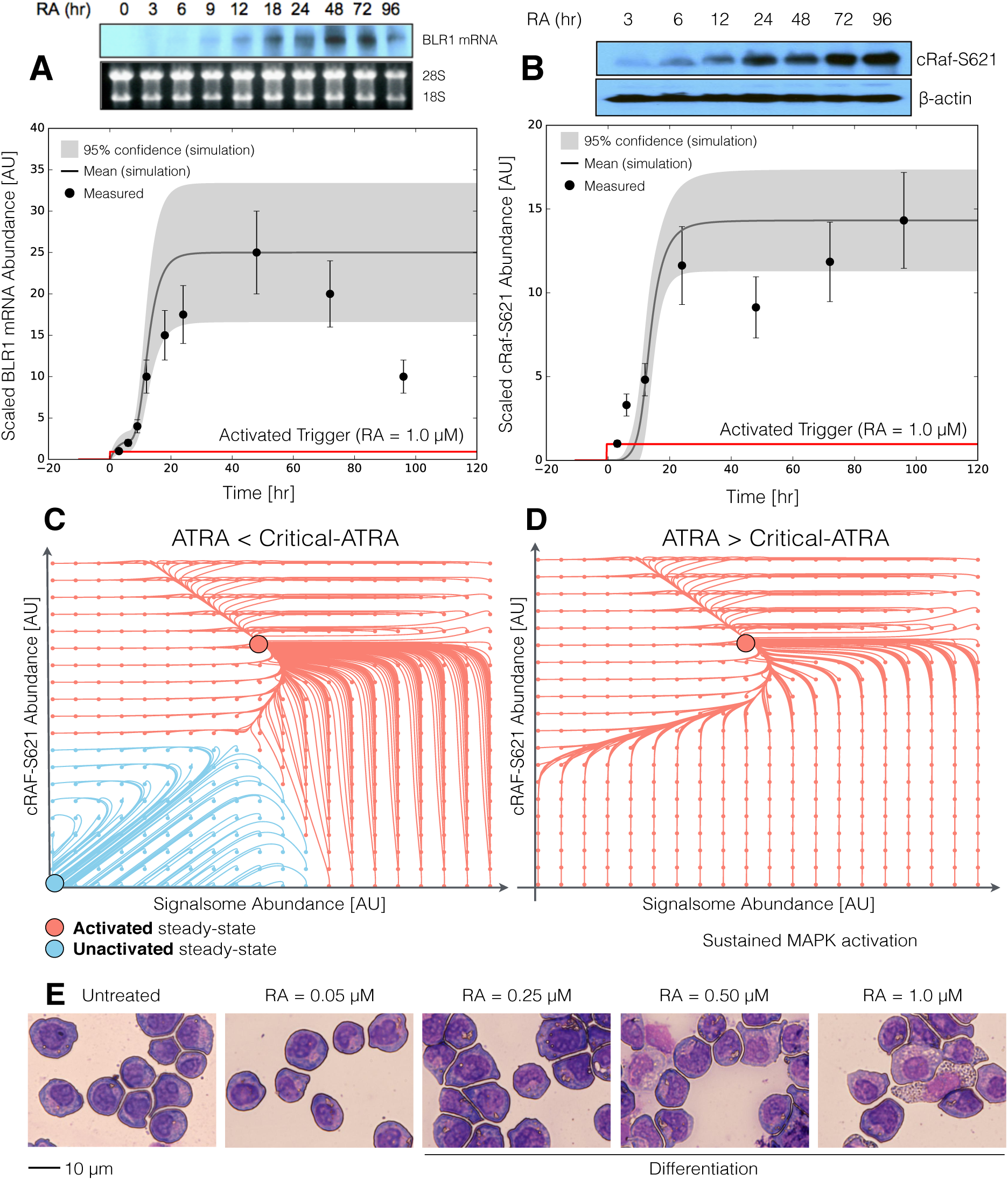
Model analysis for ATRA-induced HL-60 differentiation. A: BLR1 mRNA versus time following exposure to l*μ*M ATRA at t = 0 hr. B: cRaf-pS621 versus time following exposure to l*μ*M ATRA at t = 0 hr. Points denote experimental measurements, solid lines denote the mean model performance. Shaded regions denote the 99% confidence interval calculated over the parameter ensemble. C: Signalsome and cRaf-pS621 nullclines for ATRA below the critical threshold. The model had two stable steady states and a single unstable state in this regime. D: Signalsome and cRaf-pS621 nullclines for ATRA above the critical threshold. In this regime the model had only a single stable steady state. E: Morphology of HL-60 as a function of ATRA concentration (t = 72 hr).

The signal initiation model was bistable with respect to ATRA induction (Fig. 2C-D). Phaseplane analysis predicted two stable steady-states when ATRA was present below a critical threshold (Fig. 2C), and only a single steady-state above the threshold (Fig. 2D). In the lower stable state, neither the signalsome nor cRaf-pS621 were present (thus, the differentiation program was inactive). However, at the higher stable state, both the signalsome and cRaf-pS621 were present, allowing for sustained activation and differentiation. Interestingly, when ATRA was above a critical threshold, only the activated state was accessible (Fig. 2D). To test these findings, we first identified the ATRA threshold. We exposed HL-60 cells to different ATRA concentrations for 72 hr (Fig. 2E). Morphological changes associated with differentiation were visible for ATRA ≥ 0.25 *μ*M, suggesting the critical ATRA threshold was near this concentration. Next, we conducted ATRA washout experiments to determine if activated cells remained activated in the absence of ATRA. HL-60 cells locked into an activated state remained activated following ATRA withdraw (Fig. 3C). This sustained activation resulted from reinforcing feedback between the signalsome and the MAPK pathway. Thus, following activation, if we inhibited or removed elements from the signal initiation module we expected the signalsome and MAPK signals to decay. We simulated ATRA induced activation in the presence of kinase inhibitors, and without key circuit elements. Consistent with experimental results using multiple MAPK inhibitors, ATRA activation in the presence of MAPK inhibitors lowered the steady-state value of signalsome (Fig. 3A). In the presence of BLR1, the signalsome and cRaf-pS621 signals were maintained following ATRA withdraw (Fig. 3B, gray). On the other hand, BLR1 deletion removed the ability of the circuit to maintain a sustained MAPK response following the withdraw of ATRA (Fig. 3B, blue). Lastly, washout experiments in which cells were exposed to 1*μ*M ATRA for 24 hr, and then transferred to fresh media without ATRA, confirmed the persistence of the self sustaining activated state for up to 144 hr (Fig. 3C). Thus, these experiments confirmed that reinforcing positive feedback likely drives the ATRA-induced differentiation program. Next, we analyzed the ATRA-induced downstream gene expression program following signalsome and cRaf activation.

**Figure 3.**
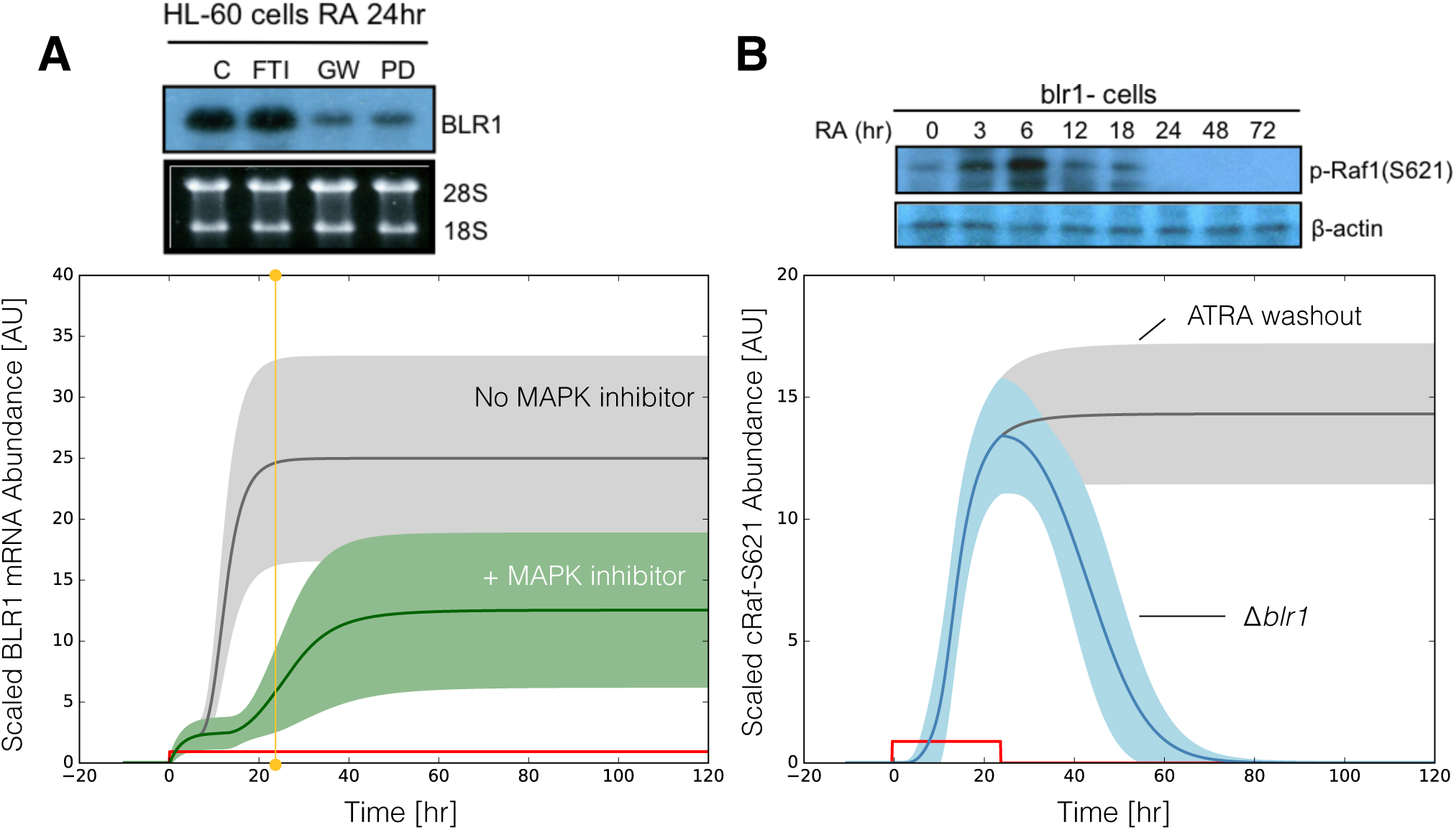
Model simulation following exposure to l*μ*M ATRA. A: BLR1 mRNA versus time with and without MAPK inhibitor. B: cRaf-pS621 versus time following pulsed exposure to l*μ*M ATRA with and without BLRl. Solid lines denote the mean model performance, while shaded regions denote the 99% confidence interval calculated over the parameter ensemble. C: Western blot analysis of phosphorylated ERKl/2 in ATRA washout experiments. Experimental data in panels A and B were reproduced from Wang and Yen^25^, data in panel Cis reported in this study.

The signal integration and phenotype modules described ATRA-induced gene expression in wild-type HL-60 cells (Fig. 4). The signal initiation module produced two outputs, activated Trigger and cRaf-pS621 which drove the expression of ATRA-induced transcription factors, which then in turn activated the phenotypic program. We assembled the connectivity of the signal integration and phenotypic programs driven by Trigger and cRaf-pS621 from literature (Table 1). We estimated the parameters for the signal initiation, and phenotype modules from steady-state and dynamic measurements of transcription factor and phenotypic marker expression following the addition of ATRA^28–31^. However, the bulk of the model parameters were taken from literature^32^ and were not estimated in this study (see materials and methods). The model simulations captured the time dependent expression of CD38 and CD11b following the addition ATRA (Fig. 4A), and the steady-state for signal integration and phenotypic markers (Fig. 4B). Lastly, we used the *predicted* values of the p21 and E2F protein abundance to estimate a blackbox model of ATRA-induced G0 arrest (Fig. 5). The phenotype module predicted p21 expression significantly increased and E2F expression decreased, in response to ATRA exposure (Fig. 5A). We then used the ratio of these values in a polynomial model to calculate the fraction of HL-60 cells in G0 arrest following the addition of ATRA (Fig. 5B). The third-order polynomial model captured the trend in measured G0-arrest values as a function of time, and was robust to uncertainty in the measured data (Fig. 5B, gray). Taken together, the output of the signal integration and phenotypic modules was consistent with time-series and steady-state measurements, thereby validating the assumed molecular connectivity. Moreover, outputs from the phenotype module described the trend in ATRA-induced G0 cell cycle arrest. Next, we explored which proteins and protein interactions in the signal integration module most influenced the system response.

**Figure 4.**
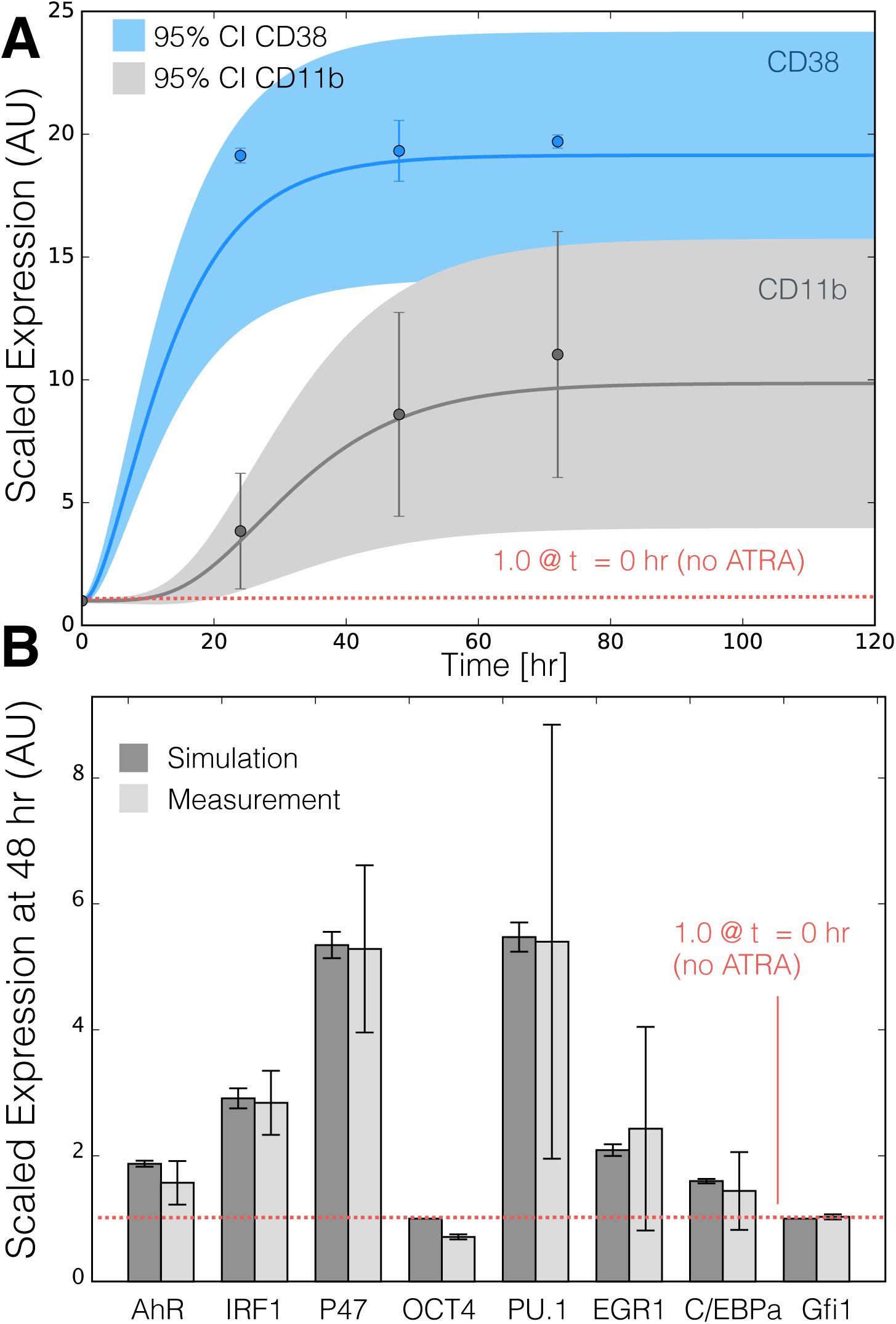
Model simulation of the HL-60 gene expression program following exposure to 1*μ*M ATRA at t = 0 hr. A: Scaled CD38 and CD11b expression versus time following ATRA exposure at time t = 0 hr. B: Scaled Gene expression at t = 48 hr following ATRA exposure. Gene expression was normalized to expression in the absence of ATRA. Experimental data in panels A and B were reproduced from Jensen et al.^31^. Model simulations were conducted using the ten best parameter sets collected during model identification. Solid lines (or bars) denote the mean model performance, while the shaded region (or error bars) denote the 95% confidence interval calculated over the parameter ensemble.

**Figure 5.**
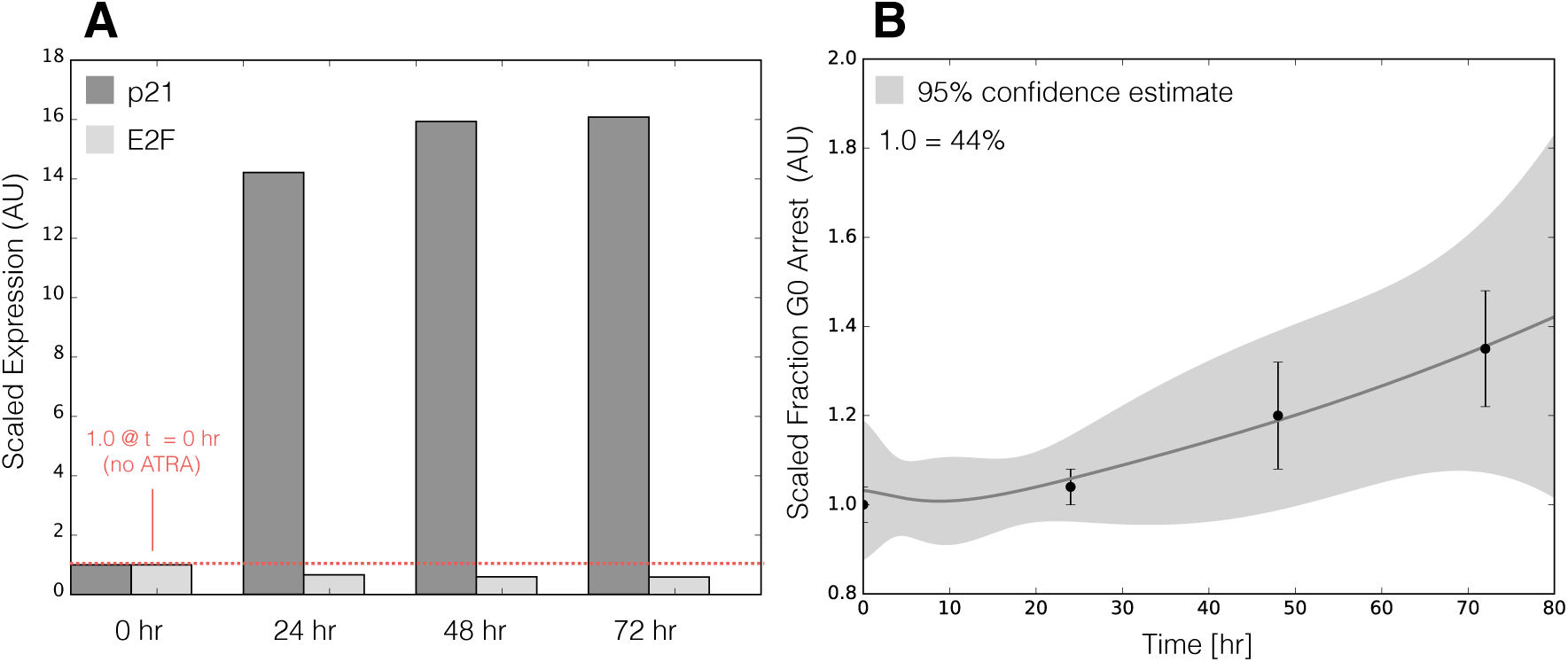
Model simulation of HL-60 cell-cycle arrest following exposure to 1*μ*M ATRA at t = 0 hr. A: Predicted p21 and E2F expression levels for the best parameter set following ATRA exposure at time t = 0 hr. B: Estimated fraction of HL-60 cells in GO arrest following ATRA exposure at time t = 0 hr. Solid lines (or bars) denote the mean model performance, while the shaded region (or error bars) denotes the 95% confidence estimate of the polynomial model. Experimental data in panel B was reproduced from Jensen et al.^31^.

The Gfi-1 and PPAR*γ* proteins were important regulators of ATRA-induced signal integration and phenotypic change (Fig. 6). We conducted pairwise gene knockout simulations in the signal integration and phenotype modules to estimate which proteins controlled the processing of the Trigger and cRaf-S621 signals. The difference between the system state with and without the gene knockouts (encoded as a normalized state displacement matrix) was decomposed using Singular Value Decomposition (SVD). A panel of ten parameter sets was sampled, and the average normalized displacement matrix was decomposed. The first six modes (approximately 36% of the total) described >95% of the gene knockout variance, with the most important components of these modes being the Gfi-1 and PPAR*γ* proteins, and to a lesser extent PU.1, C/EBP*α* and and AP1 (Fig. 6A). To better understand which protein-DNA connections were important, we simulated the pairwise deletion of interactions between these proteins and their respective regulatory targets. Singular value decomposition of the normalized state displacement matrix assembled from the pairwise connection deletions, suggested the first six modes (approximately 26% of the total) accounted for >90% of the variance. Globally, the most sensitive interactions controlled p21 and p47Phox expression, markers for cell-cycle arrest and reactive oxygen formation phenotypic axes activated following ATRA addition (Fig. 6B). Analysis of the modes suggested the action of PPAR*γ*, Gfi-1 and C/EBP*α* were consistently important over multiple target genes. The connection knockout analysis also revealed robustness in the network. For example, no pair of deletions qualitatively changed the expression of regulators such as PU.1, Oct1, Oct4 or PPAR*γ*. Thus, the expression of these species was robust to disturbance in the connectivity. To better understand the combined influence of the PPAR*γ* and Gfi-1 deletions, we computed the fold change in the protein levels in the single (Gfi-1^-/-^ or PPAR*γ*^-/-^) and double (Gfi-1^-/-^ and PPAR*γ*^-/-^) mutants for the best fit parameter set (Fig. 7). Deletion of Gfi-1 led to a 2-4 fold increase in EGR-1, CD11b and C/EBP*α* expression, and a >8 fold increase in PU.1 abundance (Fig. 7,blue). On the other hand, deletion of PPAR*γ* led to >8 fold downregulation of CD38, p21, IRF1 and Oct1 (Fig. 7,red). Both knockouts slightly increased E2F expression, but neither influenced the expression of p47Phox. The double mutant was qualitatively similar to the combined behavior of the two single mutant cases. Taken together, Gfi-1 and PPAR*γ* controlled the cell-cycle arrest and receptor signaling axes, with PPAR*γ* regulating CD38, IRF1 and p21 expression while Gfi-1 controlled CD11b expression. These simulations suggested deletion of PPAR*γ* and Gfi-1 would not interfere with reactive oxygen formation, but would limit the ability of HL-60 cells to arrest. However, this analysis did not give insight into which components upstream of the signal initiation module were important. Toward this question, we explored the composition and regulation of the signalsome complex by experimentally interrogating a panel of possible Raf interaction partners.

**Figure 6.**
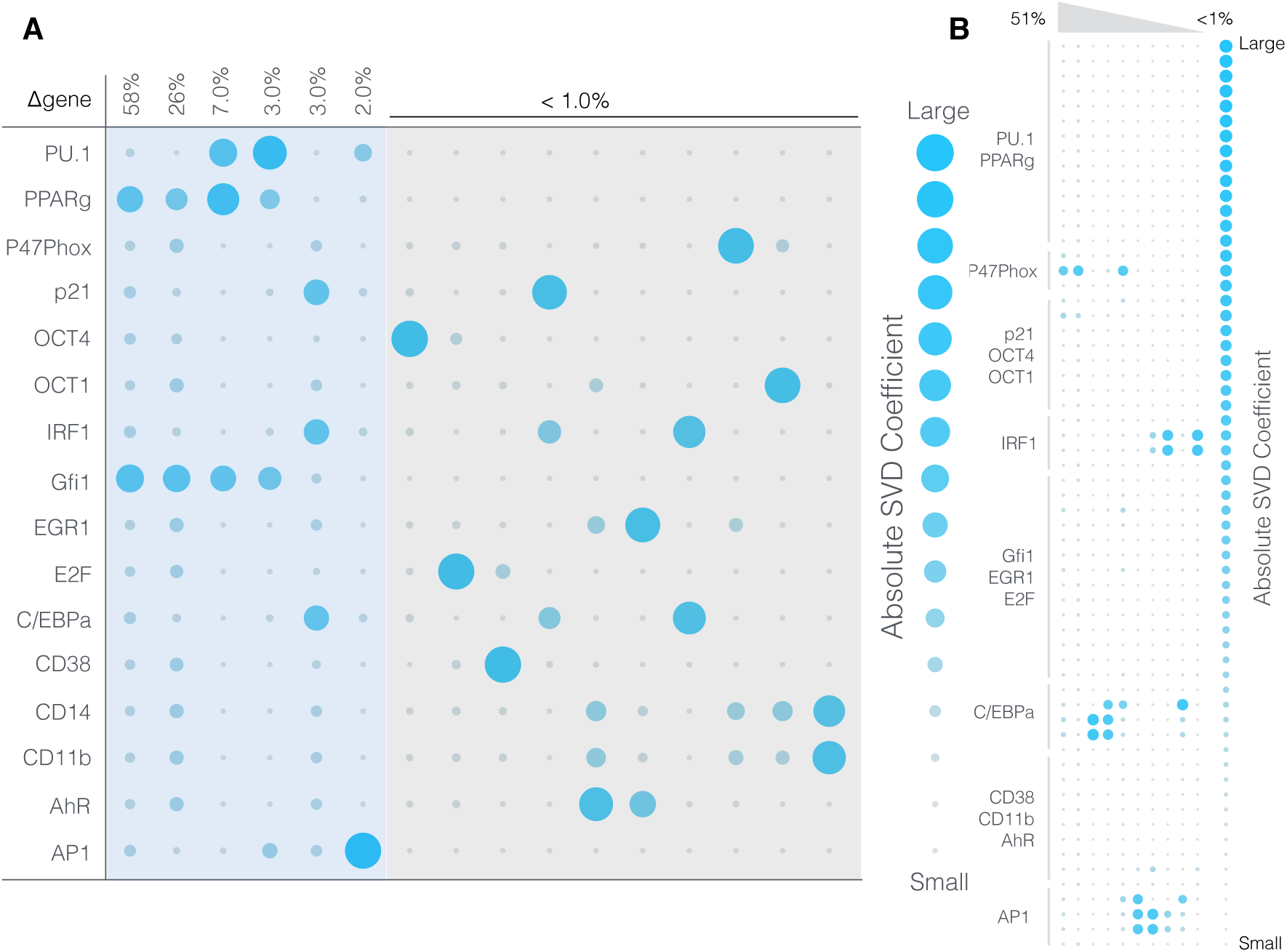
Robustness of the HL-60 differentiation program following exposure to 1*μ*M ATRA at t = 0 hr. A: Singular value decomposition of the system response (*l*^2^-norm between the perturbed and nominal state) following pairwise gene knockout simulations using the best fit parameter set. The percentage at the top of each column describes the fraction of the variance in the system state captured by the node combinations in the rows. B: Singular value decomposition of the system response (*l*^2^-norm between the perturbed and nominal state) following the pairwise removal of connections.

**Figure 7.**
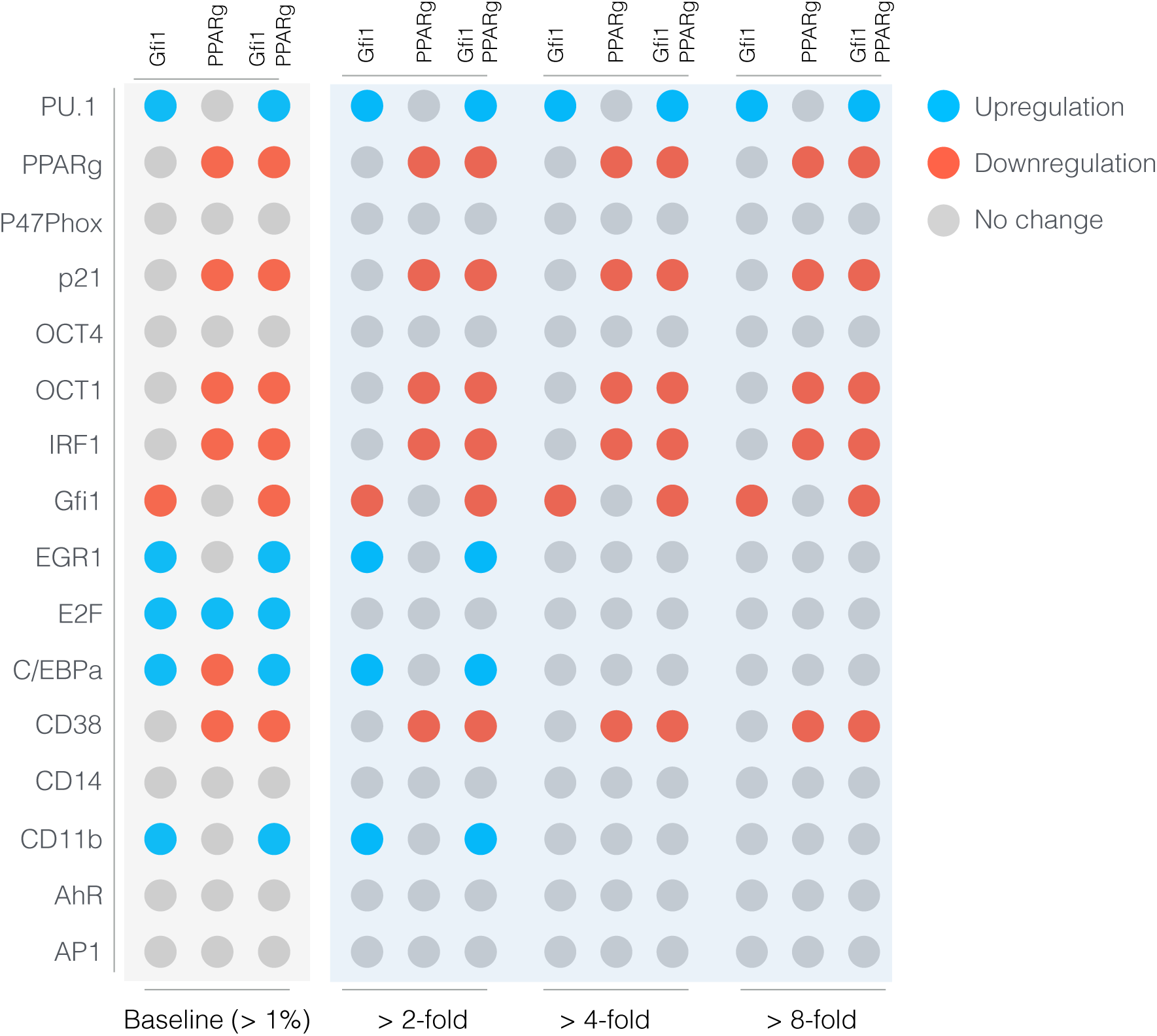
Robustness of the HL-60 differentiation program following exposure to 1*μ*M ATRA at t = 0 hr. Protein fold change at t = 48 hr (rows) in single and double knock-out mutants (columns) relative to wild-type HL-60 cells. The responses were grouped into >2,4 and 8 fold changes. The best fit parameter set was used to calculate the protein fold change.

The full composition of the signalsome, and the kinase therein ultimately responsible for mediating ATRA-induced Raf activation is still not known. To explore this question, we conducted immunoprecipitation and subsequent Western blotting to identify interactions between Raf and 19 putative interaction partners. A panel of 19 possible Raf interaction partners (kinases, GTPases, scaffolding proteins etc) was constructed based upon known signaling pathways. We did not consider the most likely binding partner, the small GTPase RAS, as previous studies have ruled it out in MAPK activation in HL-60 cells^25,33^. Total Raf was used as a bait protein for the immunoprecipitation studies. Interrogation of the Raf interactome thus suggested Vav1 was involved with ATRA-induced initiation of MAPK activity (Fig. 8). Western blot analysis using total Raf and Raf-pS621 specific antibodies confirmed the presence of the bait protein, total and phosphorylated forms, in the immunoprecipitate (Fig. 8A). Of the 19 proteins sampled, Vav1, Src, CK2, Akt, and 14-3-3 co-precipitated with Raf, suggesting their co-existence in a complex was possible. However, only the associations between Raf and Vav1, and Raf and Src were ATRA-inducible (Fig. 8). The interaction between Vav1 and Raf was one of the most prominent interactions in the panel, and it was crippled by inhibiting Raf. Furthermore, the Vav1 and Src associations were correlated with Raf-pS621 abundance in the precipitate. Other proteins e.g., CK2, Akt and 14-3-3, generally bound Raf regardless of phosphorylation status or ATRA treatment. The remaining 14 proteins were expressed in whole cell lysate (Fig. 8B), but were not detectable in the immuno-precipitate with Raf IP; consistent with the potential importance of the Raf-Vav interaction for signaling, it paralleled Raf phosphorylation at S621, a putative telltale of the activated kinase. Furthermore, treatment with the Raf kinase inhibitor GW5074 following ATRA exposure reduced the association of both Vav1 with Raf and Src with Raf (Fig. 8), although the signal intensity for Src was notably weak. However, GW5074 did not influence the association of CK2 or 14-3-3 with Raf, further demonstrating their independence from Raf phosphorylation. Interestingly, the Raf-Akt interaction qualitatively increased following treatment with GW5074; however, it remained unaffected by treatment with ATRA. Src family kinases are known to be important in myeloid differentiation^34^ and their role in HL-60 differentiation has been investigated elsewhere^15^. Given the existing work and variable reproducibility in the context of the Raf immunoprecipitate, we did not investigate the role of Src further in this study. Taken together, the immunoprecipitation and GW5074 results implicated Vav1 association to be correlated with Raf activation following ATRA-treatment. Further, while we observed possible immunoprecipitation of Src with Raf, the western blot results showed inconsistent results and significant non-specific binding; therefore we could not rule in or out a Src/Raf interaction. Previous studies demonstrated that a Vav1-Slp76-Cbl-CD38 complex plays an important role in ATRA-induced MAPK activation and differentiation of HL-60 cells^17^. Here we did not observe direct interaction of Raf with Cbl or Slp76; however, this interaction could could be involved upstream. Next, we considered the effect of the Raf kinase inhibitor GW5074 on functional markers of ATRA-induced growth arrest and differentiation.

Inhibition of Raf kinase activity modulated MAPK activation and differentiation markers following ATRA exposure (Fig. 8D-F). ATRA treatment alone statistically significantly increased the G1/G0 percentage over the untreated control, while GW5074 alone had a negligible effect on the cell cycle distribution (Fig. 8D). Surprisingly, the combination of GW5074 and ATRA statistically significantly increased the G1/G0 population (82 ± 1%) compared with ATRA alone (61 ± 0.5%). Increased G1/G0 arrest following the combined treatment with GW5074 and ATRA was unexpected, as the combination of ATRA and the MEK inhibitor (PD98059) has been shown previously to decrease ATRA-induced growth arrest^12^. However, growth arrest is not the sole indication of functional differentiation. Expression of the cell surface marker CD11b has also been shown to coincide with HL-60 cells myeloid differentiation^35^. We measured CD11b expression, for the various treatment groups, using immuno-fluorescence flow cytometry 48 hr post-treatment. As with G1/G0 arrest, ATRA alone increased CD11b expression over the untreated control, while GW5074 further enhanced ATRA-induced CD11b expression (Fig. 8E). GW5074 alone had no statistically significant effect on CD11b expression, compared with the untreated control. Lastly, the inducible reactive oxygen species (ROS) response was used as a functional marker of differentiated neutrophils^20^. We measured the ROS response induced by the phorbol ester 12-O-tetradecanoylphorbol-13-acetate (TPA) using flow cytometry. Untreated cells showed no discernible TPA response, with only 7.0 ± 3.0% ROS induction (Fig. 8F). Cells treated with ATRA had a significantly increased TPA response, 53 ± 7% ROS induction 48 hr post-treatment. Treatment with both ATRA and GW5074 statistically significantly reduced ROS induction (22 ± 0.6%) compared to ATRA alone. Interestingly, Western blot analysis did not detect a GW5074 effect on ATRA-induced expression of p47Phox, a required upstream component of the ROS response (Fig. 8F, bottom). Thus, the inhibitory effect of GW5074 on inducible ROS might occur downstream of p47Phox expression. However, the ROS producing complex is MAPK dependent, therefore it is also possible that GW5074 inhibited ROS production by interfering with MAPK activation (in which case the p47Phox marker might not accurately reflect phenotypic conversion and differentiation).

**Figure 8.**
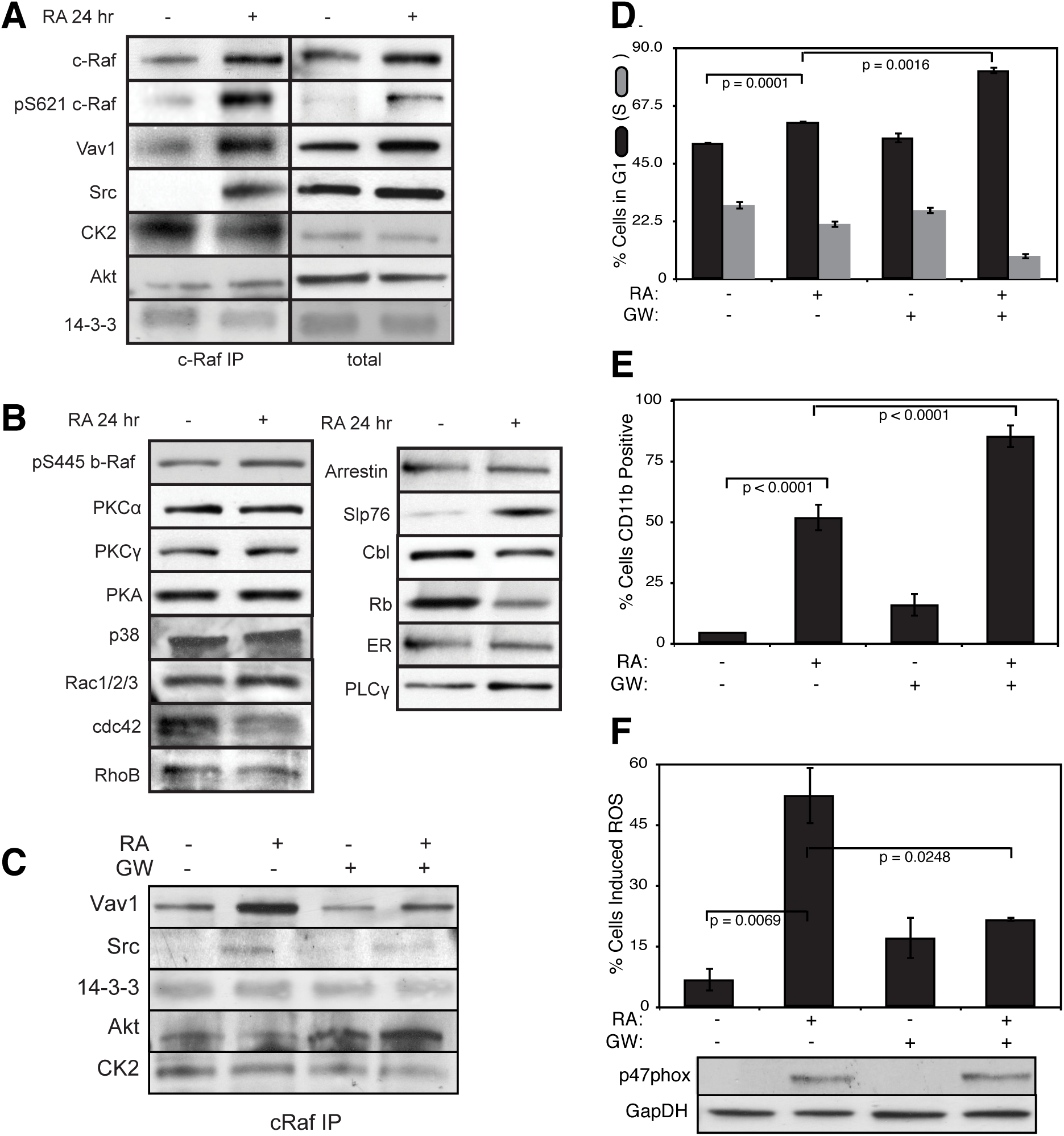
Investigation of a panel of possible Raf interaction partners in the presence and absence of ATRA. A: Species identified to precipitate out with Raf: first column shows Western blot analysis on total Raf immunoprecipitation with and without 24 hr ATRA treatment and the second on total lysate. B: The expression of species considered that did not precipitate out with Raf at levels detectable by Western blot analysis on total lysate. C: Effect of the Raf inhibitor GW5074 on Raf interactions as determined by Western blot analysis of total Raf immunoprecipitation. The Authors note the the signal associated with Src was weak. D: Cell Cycle distribution as determined by flow cytometry indicated arrest induced by ATRA, which was increased by the addition of GW5074. E: Expression of the cell surface marker CD11b as determined by flow cytometry indicated increased expression induced by ATRA, which was enhanced by the addition of GW5074. F: Inducible reactive oxygen species (ROS) as determined by DCF flow cytometry. The functional differentiation response of ATRA treated cells was mitigated by GW5074.

## Discussion

In this study, we presented an effective model of ATRA-inducible differentiation of HL-60 cells. The model consisted of three modules: a signal initiation module that sensed and transformed the ATRA signal into activated cRaf-pS621 and the ATRA-RAR/RXR (Trigger) signals; a signal integration module that controlled the expression of upstream transcription factors given cRaf-pS621 and activated Trigger signals; and a phenotype module which encoded the expression of functional differentiation markers from the ATRA-inducible transcription factors. The model described the transcription and translation of genes in each module, and signaling events in each module in a growing population of HL-60 cells. Model parameters were taken from literature, however, unknown coefficients that appear in the promoter logic models were estimated from protein measurements in HL-60 cells following ATRA exposure. Despite its simplicity, the effective model captured key features of the ATRA induced differentiation such as sustained MAPK activation, and bistability with respect to ATRA exposure. The model also described the expression of upstream transcription factors which regulated the expression of differentiation markers. Lastly, analysis of the response of the model to perturbations identified Gfi-1 and PPAR*γ* as master regulators of ATRA-induced differentiation. We also found evidence of a prominent regulatory role for a signaling molecule ATRA-inducible component of the signalsome, Vav1. Vav1 is a guanine nucleotide exchange factor for Rho family GTPases that activate pathways leading to actin cytoskeletal rearrangements and transcriptional alterations^36^. The Vav1/Raf association correlated with Raf activity, was ATRA-inducible and decreased after treatment with the Raf inhibitor GW5074.

Naturally occurring cell fate decisions often incorporate reinforcing feedback and bistability^37,38^. One of the most well studied cell fate circuits is the Mos mitogen-activated protein kinase cascade in *Xenopus* oocytes. This cascade is activated when oocytes are induced by the steroid hormone progesterone^39^. The MEK-dependent activation of p42 MAPK stimulates the accumulation of the Mos oncoprotein, which in turn activates MEK, thereby closing the feedback loop. This is similar to the signal initiation module presented here; ATRA drives signalsome formation, which activates MAPK, which in turn leads to more signalsome activation. Thus, while HL-60 and *Xenopus* oocytes are vastly different biological models, their cell fate programs share a similar architectural feature. Reinforcing feedback and bistability has also been implicated in hematopoietic cell fate determination. Laslo et al showed in nonmalignant myelomonocytic cells that the counter antagonistic repressors, Gfi-1 and Egr-1/2 (whose expression is tuned by PU.1 and C/EBP*α*), encode a bistable switch that results in a macrophage, neutrophil or a mixed lineage population depending upon PU.1 and C/EBP*α* expression^38^. The current model contained the Gfi-1 and Egr-1/2 agonistic switch; however, its significance was unclear for HL-60 cells. The expression of Gfi-1, Egr-1/2, C/EBP*α* and PU.1 was not consistent with the canonical lineage pattern expected from literature. For example, Egr-1/2 expression (associated with a macrophage lineage) increased, while Gfi-1 expression (associated with a neutrophil lineage) was unchanged following ATRA exposure. Thus, HL-60 cells, which are a less mature cancer cell line, exhibited a non-canonical expression pattern. Other unrelated cell fate decisions such as programmed cell death have also been suggested to be bistable^40^. Still more biochemical networks important to human health, for example the human coagulation or complement cascades, also feature strong positive feedback elements^41^. Thus, while reinforcing feedback is often undesirable in human engineered systems, it is at the core of a diverse variety of cell fate programs and other networks important to human health.

Analysis of the signal integration and phenotype modules suggested Gfi-1 and PPAR*γ* proteins were important regulators of ATRA-induced signal integration and phenotypic change. Model analysis showed that PU.1, Egr-1 and C/EBP*α* expression increased in Gfi-1^-/-^ mutants, where PU.1 expression was upregulated by greater than 8-fold. PU.1, a member of the *ets* transcription factor family, is a well known regulator of granulocyte and monocyte development^42^. The relative level of PU.1 and C/EBP*α* is thought to control macrophage versus neutrophil cell fate decisions in granulocytic macrophage progenitor cells^43^. Simulations suggested that combined Gfi-1 + PPAR*γ* deletion crippled the ability of HL-60 cells to undergo neutrophilic differentiation following ATRA exposure. p21 expression decreased significantly, suggesting Gfi-1^-/-^ + PPAR*γ*^-/-^ mutants were less likely to G0-arrest following ATRA exposure. The expression of other neutrophilic markers, such as CD38, also decreased in Gfi-1^-/-^ + PPAR*γ*^-/-^ cells. On the other hand, the expression of reactive oxygen metabolic markers, or other important transcription factors such as Oct4 did not change. For example, model analysis suggested that the C/EBP*α* dependent interaction of PU.1 with the *NCF1* gene, which encodes the p47Phox protein, was the most sensitive PU.1 connection; deletion of this connection removed the ability of the system to express p47Phox. p47Phox, also known as neutrophil cytosol factor 1, is one of four cytosolic subunits of the multi-protein NADPH oxidase complex found in neutrophils^44^. This enzyme is responsible for reactive oxygen species (ROS) production, a key component of the anti-microbial function of neutrophils. While p47Phox expression required C/EBP*α* and PU.1, neither Gfi-1 nor PPAR*γ* deletion increased expression. This suggested that p47Phox expression was saturated with respect to C/EBP*α* and PU.1, and simultaneously not sensitive to PPAR*γ* abundance. Taken together, Gfi-1^-/-^ + PPAR*γ*^-/-^ cells were predicted to exhibit some aspects of the ATRA response, but not other critical features such as cell cycle arrest. Hock et al showed that Gfi-1^-/-^ mice lacked normal neutrophils, and were highly sensitive to bacterial infection^45^. Thus, the model analysis was consistent with this study. However, other predictions concerning the behavior of the Gfi-1^-/-^ + PPAR*γ*^-/-^ mutants remain to be tested.

Immunoprecipitation studies identified a limited number of ATRA-dependent and -independent Raf interaction partners. While we were unable to detect the association of Raf with common kinases and GTPases such as PKC, PKA, p38, Rac and Rho, we did establish potential interactions between Raf and key partners such as Vav1, Src, Akt, CK2 and 14-3-3. All of these partners are known to be associated with Raf activation or function. Src is known to bind Raf through an SH2 domain, and this association has been shown to be dependent of the serine phosphorylation of Raf^46^. Thus, an ATRA inducible Src/Raf association may be a result of ATRA-induced Raf phosphorylation at S259 or S621. We also identified an interaction between Raf and the Ser/Thr kinases Akt and CK2. Akt can phosphorylate Raf at S259, as demonstrated by studies in a human breast cancer line^47^. CK2 can also phosphorylate Raf, although the literature has traditionally focused on S338 and not S621 or S259^48^. However, neither of these kinase interactions were ATRA-inducible, suggesting their association with Raf alone was not associated with ATRA-induced Raf phosphorylation. The adapter protein 14-3-3 was also constitutively associated with Raf. The interaction between Raf and 14-3-3 has been associated with both S621 and S259 phosphorylation and activity^49^. Additionally, the association of Raf with 14-3-3 not only stabilized S621 phosphorylation, but also reversed the S621 phosphorylation from inhibitory to activating^50^. Finally, we found that Vav1/Raf association correlated with Raf activity, was ATRA-inducible and decreased after treatment with GW5074. The presence of Vav1 in Raf/Grb2 complexes has been shown to correlate with increased Raf activity in mast cells^51^. Furthermore, studies on Vav1 knockout mice demonstrated that the loss of Vav1 resulted in deficiencies of ERK signaling for both T-cells as well as neutrophils^52,53^. Interestingly, while an integrin ligand-induced ROS response was blocked in Vav1 knockout neutrophils, TPA was able to bypass the Vav1 requirement and stimulate both ERK phosphorylation and ROS induction^53^. In this study, the TPA-induced ROS response was dependent upon Raf kinase activity, and was mitigated by the addition of GW5074. It is possible that Vav1 is downstream of various integrin receptors but upstream of Raf in terms of inducible ROS responses. Vav1 has also been shown to associate with a Cbl-Slp76-CD38 complex in an ATRA-dependent manner; furthermore, transfection of HL-60 cells with Cbl mutants that fail to bind CD38, yet still bind Slp76 and Vav1, prevents ATRA-induced MAPK activation^17^. The literature suggest a variety of possible receptor-signaling pathways, which involve Vav1, for MAPK activation; moreover, given the ATRA-inducible association Vav1 may play a direct role in Raf activation.

We hypothesized that Vav1 is a member of an ATRA-inducible signalsome complex which propels sustained MAPK activation, arrest and differentiation (shown schematically in Fig. 9). Initially, ATRA-induced Vav1 expression drives increased association between Vav1 and Raf. This increased interaction facilitates phosphorylation and activation of Raf by pre-bound Akt and/or CK2 at S621 or perhaps S259. Constitutively bound 14-3-3 may also stabilize the S621 phosphorylation, modulate the activity and/or up-regulate autophosphorylation. Activated Raf can then drive ERK activation, which in turn closes the positive feedback loop by activating Raf transcription factors e.g., Sp1 and/or STAT1^54–57^. We tested this working hypothesis using mathematical modeling. The model recapitulated both ATRA time-course data as well as the GW5074 inhibitor effects. This suggested the proposed Raf-Vav1 architecture was at least consistent with the experimental studies. Further, analysis of the Raf-Vav1 model identified bistability in phosphorylated ERK levels. Thus, two possible MAPK activation branches were possible for experimentally testable ATRA values. The analysis also suggested the ATRA-induced Raf-Vav1 architecture could be locked into a sustained signaling mode (high phosphorylated ERK) even in the absence of a ATRA signal. This locked-in property could give rise to an ATRA-induction memory. We validated the treatment memory property predicted by the Raf-Vav1 circuit experimentally using ATRA-washout experiments. ERK phosphorylation levels remained high for more then 96 hr after ATRA was removed. Previous studies demonstrated that HL-60 cells possessed an inheritable memory of ATRA stimulus^58^. Although the active state was self-sustaining, the inactive state demonstrated considerable robustness to perturbation. For example, we found that 50x overexpression of Raf was required to reliably lock MAPK into the activated state, while small perturbations had almost no effect on phosphorylated ERK levels over the entire ensemble. CD38 expression correlated with the phosphorylated ERK, suggesting its involvement in the signaling complex. Our computational and experimental results showed that positive feedback, through ERK-dependent Raf expression, could sustain MAPK signaling through many division cycles. Such molecular mechanisms could underly aspects of cellular memory associated to consecutive ATRA treatments.

**Figure 9.**
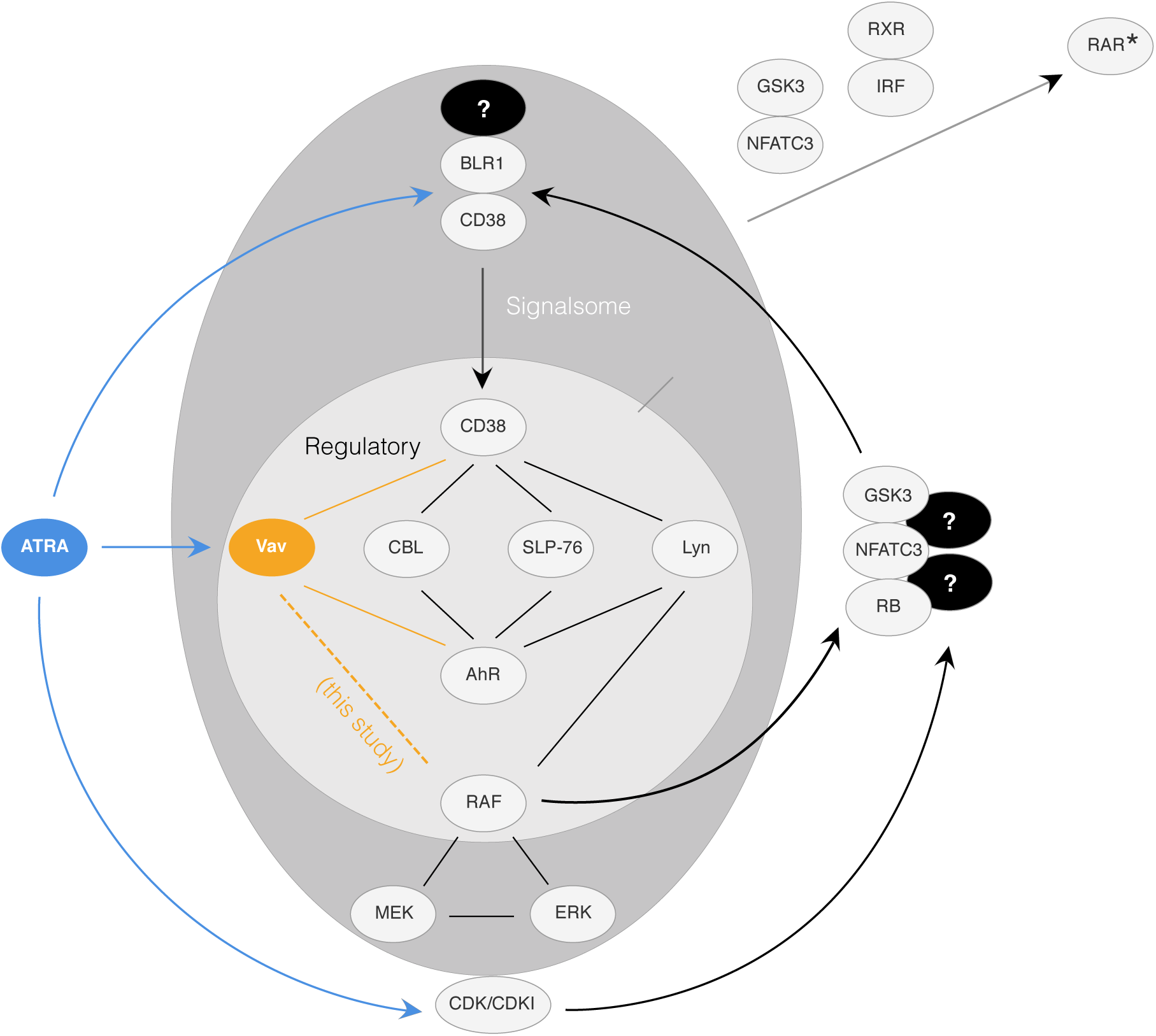
This schematic diagram shows the hypothetical principal pathways in the ATRA-induced signaling that results in cell differentiation in the HL-60 myeloid leukemia model^17, 65–69^, It is based on modules and feedback loops. There are three main arms (shown top to bottom): 1. Direct RA targeting of RAREs in genes such as CD38 or BLR1; 2. Formation of a signalsome that has a regulatory module that includes Vav (a guanine nucleotide exchange factor), CBL and SLP-76 (adaptors), and Lyn (a Src family kinase) that regulates a Raf/Mek/Erk axis that incorporates Erk to Raf feedback, where the regulators are modulated by AhR and CD38 receptors; and 3. Direct RA targeted up regulation of CDKI to control RB hypophosphorylation. The Raf/Mek/Erk axis is embedded in the signalsome and subject to modulation by the regulators. The output of the signalsome is discharge of the Raf from the cytosol to the nucleus where it binds (hyper)phospho-RB and other targets, including NFATc3, which enables activation of the RA bound RAR/RXR poised on the BLR1 promoter, and also GSK3, phosphorylation of which relieves its inhibitory effect on RAR*α*. CDKI directed hypophosphorylation of RB releases Raf sequestered by RB to go to NFATc3, GSK3, and other targets. A significant consequence of the nuclear RAF is ergo ultimately to enable or hyperactivate transcriptional activation by RAR*α* to drive differentiation. It might be noted that this proposed general model provides a mechanistic rationalization for why cell cycle arrest is historically oft times perceived as a precondition for phenotypic maturation.

## Methods

### Effective gene expression model equations

The ATRA differentiation model was encoded as a system of differential algebraic equations (DAEs) which described both signaling and gene expression processes. We modeled transcription and translation as Ordinary Differential Equations (ODEs), while signaling processes were assumed to quickly equilibrate and were treated as a pseudo steady state system of algebraic equations. The model formulation follows from a previous study of the Epithelial Messecnchymal Transition (EMT)^59^; in the current study additional attention was paid to the formulation of the transcription and translation rates, and an updated approach was taken to model the regulation of gene expression.

We decomposed the ATRA-induced differentiation program into three modules; a signal initiation module that sensed and transformed the ATRA signal into activated cRaf-pS621 and the ATRA-RAR/RXR (activated Trigger) signals; a signal integration module that controlled the expression of upstream transcription factors given cRaf-pS621 and activated Trigger signals; and a phenotype module which encoded the expression of functional differentiation markers from the ATRA-inducible transcription factors. The output of the signal initiation module was the input to the gene expression model. For each gene 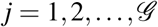, we modeled both the mRNA (*m*_*j*_), protein (*p*_*j*_) and signaling species abundance:

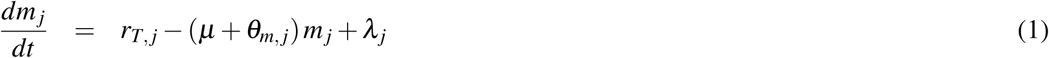

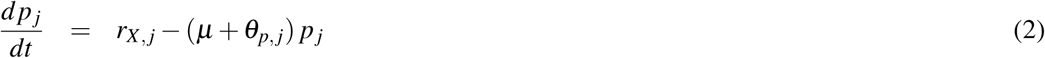

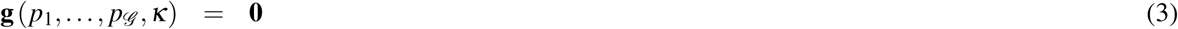

where signaling species abundance was governed by the non-linear algebraic equations 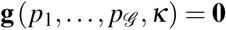. The model parameter vector is denoted by ***κ***. The terms *r*_*T, j*_ and *r*_*X, j*_ denote the specific rates of transcription, and translation while the terms *θ*_*m, j*_ and *θ*_*p, j*_ denote first-order degradation constants for mRNA and protein, respectively. The specific transcription rate *r*_*T, j*_ was modeled as the product of a kinetic term 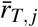 and a control term *u*_*j*_ which described how the abundance of transcription factors, or other regulators influenced the expression of gene *j*.

The gene expression control term 0 ≤ *u*_*j*_ ≤ 1 depended upon the combination of factors which influenced the expression of gene *j*. If the expression of gene *j* was influenced by 1, …, *m* factors, we modeled this relationship as 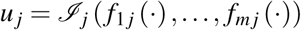 where 0 ≤ *f*_*ij*_ (⋅) ≤ 1 denotes a regulatory transfer function quantifying the influence of factor *i* on the expression of gene *j*, and 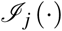 denotes an integration rule which combines the individual regulatory inputs for gene *j* into a single control term. In this study, the integration rule governing gene expression was the weighted fraction of promoter configurations that resulted in gene expression^60^:

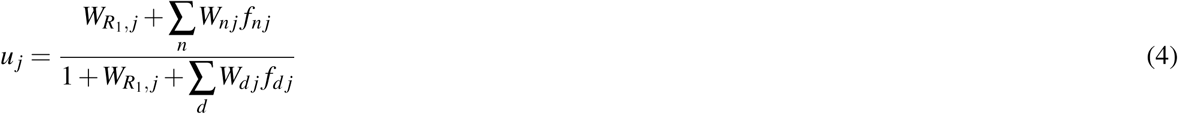

The numerator, the weighted sum (with weights *W*_*nj*_) of promoter configurations leading to gene expression, was normalized by all possible promoter configurations (denominator). The likelihood of each configuration was quantified by the transfer function *f*_*nj*_ (which we modeled using Hill functions), while the lead term in the numerator *W*_*R*1, *j*_ denotes the weight of constitutive expression for gene *j*. Given the formulation of the control law, the *λ*_*j*_ term (which denotes the constitutive rate of expression of gene *j*) was given by:

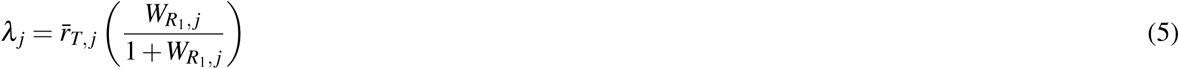

The kinetic transcription term 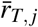 was modeled as:

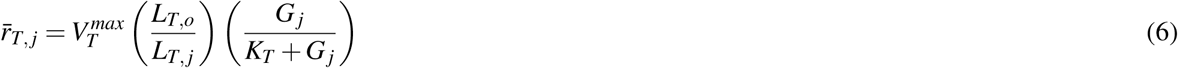

where the maximum gene expression rate 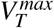 was defined as the product of a characteristic transcription rate constant (*k*_*T*_) and the abundance of RNA polymerase (*R*_1_), 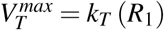. The (*L*_*T*, *o*_/*L*_*T, j*_) term denotes the ratio of transcription read lengths; *L*_*T, o*_ represents a characteristic gene length, while *L*_*T, j*_ denotes the length of gene *j*. Thus, the ratio (*L*_*T, o*_/*L*_*T, j*_) is a gene specific correction to the characteristic transcription rate 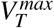. If a gene expression process had no modifying factors, *u*_*j*_ = 1. Lastly, the specific translation rate was modeled as:

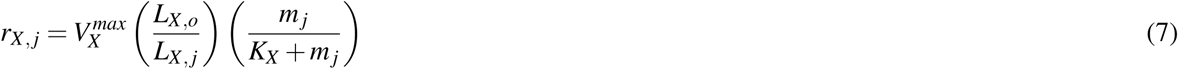

where 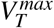 denotes a characteristic maximum translation rate estimated from literature, and *K*_*X*_ denotes a translation saturation constant. The characteristic maximum translation rate was defined as the product of a characteristic translation rate constant (*k*_*X*_) and the Ribosome abundance (*R*_2_), 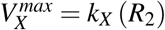. As was the case for transcription, we corrected the characteristic translation rate by the ratio of the length of a characteristic transcript normalized by the length of transcript *j*. The sequence lengths used in this study are given in Table 3; the characteristic gene and mRNA lengths were given by the average lengths computed from the values in Table 3.

**Table 3.** Sequence lengths from NCBI RefSeq database were used in the signal integration and phenotype modules^126^. The RNA sequence length used represents the total distance of transcription, and assume to be equal to the gene length.

| Gene Name | Gene (bp) | RNA (bp) | Protein (aa) | Gene ID | Protein ID |
| --- | --- | --- | --- | --- | --- |
| AP-1 | 10323 | 10323 | 331 | Gene ID: 3725 | NP_002219 |
| AhR | 47530 | 47530 | 848 | Gene ID: 196 | NP_001621 |
| CD11b | 72925 | 72925 | 1153 | Gene ID: 3684 | NP_001139280 |
| CD14 | 8974 | 8974 | 375 | Gene ID: 929 | NP_001035110 |
| CD38 | 174978 | 74978 | 300 | Gene ID: 952 | NP_001766 |
| C/EBPα | 2630 | 2630 | 393 | Gene ID: 1050 | NP_001274353.1 |
| E2F | 17919 | 17919 | 437 | Gene ID: 1869 | NP_005216 |
| Egr-1 | 10824 | 10824 | 543 | Gene ID: 1958 | NP_001955 |
| Gfi-1 | 13833 | 13833 | 422 | Gene ID: 2672 | NP_005254 |
| IRF-1 | 16165 | 16165 | 325 | Gene ID: 3659 | NP_002189 |
| Oct1 | 206516 | 206516 | 741.33 | Gene ID: 5451 | NP_002688.3, NP_001185712.1, NP_001185715.1 |
| Oct4 | 6356 | 6356 | 206.33 | Gene ID: 5460 | NP_001167002, NP_001167015, NP_001167016 |
| P21 | 15651 | 15651 | 198 | NG_009364.1 | NP_001621 |
| P47 | 3074 | 3074 | 390 | GenBank: AF003533.1 | NP_000256 |
| PPARγ | 153507 | 153507 | 250 | Gene ID: 5468 | NP_001317544 |
| PU.1 | 40782 | 40782 | 270.5 | Gene ID: 6688 | NP_001074016, NP_003111 |

### Signaling model equations

The signal initiation and integration modules required the abundance of cRaf-pS621 and ATRA-RAR/RXR (activated Trigger) as inputs. However, the base model described only the abundance of inactive proteins e.g., cRaf or RAR/RXR but not the activated forms. To address this issue, we estimated pseudo steady state approximations for the abundance of cRaf-pS621 and activated Trigger. The abundance of activated trigger (*x*_*a*,1_) was estimated directly from the RAR/RXR abundance (*x*_*u*,1_):

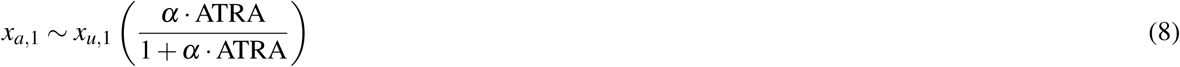

where *α* denotes a gain parameter; *α* = 0.0 if ATRA is less than a threshold, and *α* = 0.1 if ATRA is greater than the differentiation threshold. The abundance of cRaf-pS621 was estimated by making the pseudo steady state approximation on the cRaf-pS621 balance. In general, the abundance of an activated signaling species *i* was governed by:

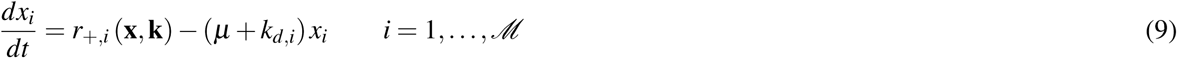

The quantity *x*_*i*_ denotes concentration of signaling species *i*, while 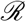 and 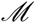 denote the number of signaling reactions and signaling species in the model, respectively. The term *r*_+,*i*_ (**x**, **k**) denotes the rate of generation of activated species *i*, while *μ* denotes the specific growth rate, and *k*_*d,i*_ denotes the rate constant controlling the non-specific degradation of *x*_*i*_. We neglected deactivation reactions e.g., phosphatase activities. We assumed that signaling processes were fast compared to gene expression; this allowed us to approximate the signaling balance as:

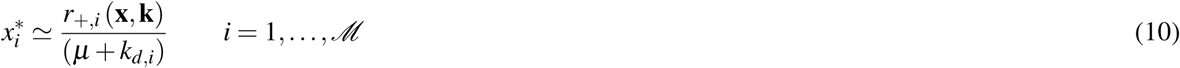

The generation rate was written as the product of a kinetic term 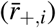 and a control term (*v*_*i*_). The control terms 0 ≤ *v*_*j*_ ≤ 1 depended upon the combination of factors which influenced rate process *j*. If rate *j* was influenced by 1, …, *m* factors, we modeled this relationship as 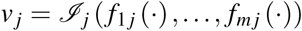 where 0 ≤ *f*_*ij*_ (⋅) ≤ 1 denotes a regulatory transfer function quantifying the influence of factor *i* on rate *j*. The function 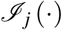 is an integration rule which maps the output of regulatory transfer functions into a control variable. In this study, we used 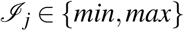 and hill transfer functions^61^. If a process had no modifying factors, *v*_*j*_ = 1. The kinetic rate of cRaf-pS621 generation 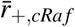 was modeled as:

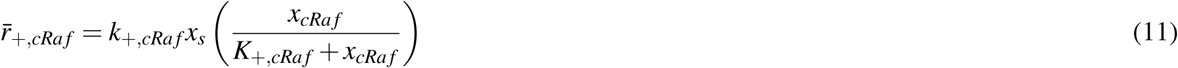

where *x*_*s*_ denotes the signalsome abundance, *k*_+,*cRaf*_ denotes a characteristic activation rate constant, and *K*_+,*cRaf*_ denotes a saturation constant governing cRaf-pS621 formation. In thus study, signalsome abundance was approximated by the abundance of the BLR1 protein; BLR1 expression is directly related to Raf nuclear translocation which in turn is related to activated signalsome. Thus, BLR1 is an indirect measure of the signalsome. The formation of cRaf-pS621 was regulated by only a single factor, the abundance of MAPK inhibitor, thus *v*_+,*cRaf*_ took the form:

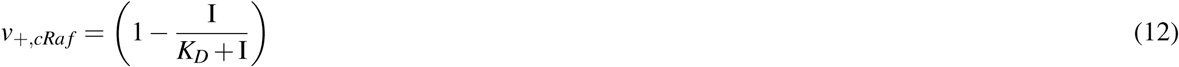

where I denotes the abundance of the MAPK inhibitor, and *K*_*D*_ denotes the inhibitor affinity.

### Estimation of gene expression model parameters

Parameters appearing in the mRNA and protein balances, e.g., maximum transcription and translation rates, the half-life of a typical mRNA and proteins (assumed to be same for all transcripts/proteins), and typical values for the copies per cell of RNA polymerase and ribosomes were estimated from literature (Table 2). The saturation constants *K*_*X*_ and *K*_*T*_ appearing in the transcription and translation rate equations were adjusted so that gene expression and translation resulted in gene products on a biologically realistic concentration scale. Lastly, we calculated the concentration for gene *G*_*j*_ by assuming, on average, that a cell had two copies of each gene at any given time. Thus, the bulk of our model parameters were taken from literature, and were not adjusted during model identification. However, the remaining parameters, e.g., the *W*_*ij*_ values or parameters appearing in the transfer functions *f*_*dj*_ which appeared in the gene expression control laws, were estimated from the experimental data discussed here. We assumed promoter configuration weights were bounded between *W*_*ij*_ ∈ [0, 100]; all cooperativity coefficients ***η***_*ij*_ appearing in the binding transfer functions *f*_*dj*_ were bounded between ***η***_*ij*_ ∈ [0, 4]; and all disassociation constants *K*_*ij*_ appearing in the binding transfer functions *f*_*dj*_ were bounded between *K*_*ij*_ ∈ [0, 1000] (nM).

Signaling and gene expression model parameters were estimated by minimizing the squared difference between simulations and experimental protein data set *j*. We measured the squared difference in the scale, fold change and shape for protein *j*:

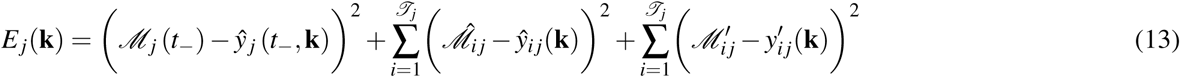

The first term in Eq. (13) quantified the initial *scale* error, directly before the addition of ATRA. In this case, 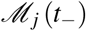 (the approximate concentration of protein *j* before the addition of ATRA) was estimated from literature. This term was required because the protein measurements were reported as the fold-change; thus, the data was normalized by a control value measured before the addition of ATRA. However, the model operated on a physical scale. The first term allowed the model to capture physically realistic changes following ATRA addition. The second term quantified the difference in the *fold-change* of protein *j* as a function of time. The terms 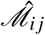 and 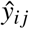 denote the scaled experimental observations and simulation outputs (fold-change; protein normalized by control value directly before ATRA addition) at time *i* from protein *j*, where 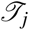 denoted the number of time points for data set *j*. Lastly, the third term of the objective function measured the difference in the *shape* of the measured and simulated protein levels. The scaled value 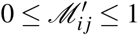 was given by:

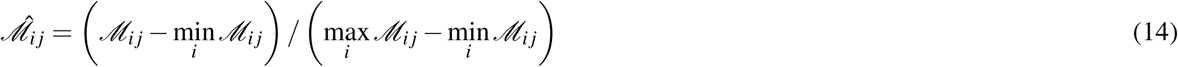

where 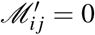 and 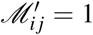 describe the lowest (highest) intensity bands. A similar scaling was used for the simulation output. We minimized the total model residual Σ_*j*_*E*_*j*_ using a heuristic direct-search optimization procedure, subject to box constraints on the parameter values, starting from a random initial parameter guess. Each downhill step was archived and used for ensemble calculations. The optimization procedure (a covariance matrix adaptation evolution strategy) has been reported previously^62^.

### Estimation of an effective cell cycle arrest model

We formulated an effective N-order polynomial model of the fraction of cells undergoing ATRA-induced cell cycle arrest at time t, 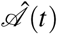, as:

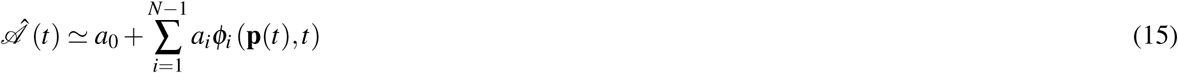

where *a*_*i*_ were unknown parameters, and *ϕ*_*i*_ (**p**(*t*), *t*) denotes a basis function. The basis functions were dependent upon the system state; in this study, we used N = 4 and basis functions of the form:

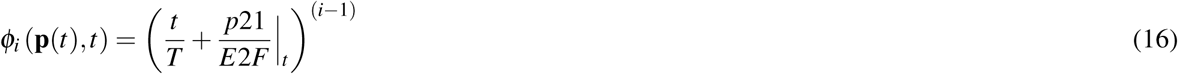

The parameters *a*_0_, …, *a*_3_ were estimated directly from cell-cycle measurements (biological replicates) using least-squares. The form of the basis function assumed p21 was directly proportional, and E2F inversely proportional, to G0-arrest. However, this was one of many possible forms for the basis functions.

### Cell culture and treatment

Human myeloblastic leukemia cells (HL-60 cells) were grown in a humidified atmosphere of 5% CO_2_ at 37*°*C and maintained in RPMI 1640 from Gibco (Carlsbad, CA) supplemented with 5% heat inactivated fetal bovine serum from Hyclone (Logan, UT) and 1 × antibiotic/antimicotic (Gibco, Carlsbad, CA). Cells were cultured in constant exponential growth^63^. Experimental cultures were initiated at 0.1 × 10^6^ cells/mL 24 hr prior to ATRA treatment; if indicated, cells were also treated with GW5074 (2*μ*M) 18 hr before ATRA treatment. For the cell culture washout experiments, cells were treated with ATRA for 24 hr, washed 3x with prewarmed serum supplemented culture medium to remove ATRA, and reseeded in ATRA-free media as described. Western blot analysis was performed at incremental time points after removal of ATRA.

### Chemicals

All-Trans Retinoic Acid (ATRA) from Sigma-Aldrich (St. Louis, MO) was dissolved in 100% ethanol with a stock concentration of 5mM, and used at a final concentration of 1*μ*M (unless otherwise noted). The cRaf inhibitor GW5074 from Sigma-Aldrich (St. Louis, MO) was dissolved in DMSO with a stock concentration of 10mM, and used at a final concentration of 2*μ*M. HL-60 cells were treated with 2*μ*M GW5074 with or without ATRA (1*μ*M) at 0 hr. This GW5074 dosage had a negligible effect on the cell cycle distribution, compared to ATRA treatment alone.

### Immunoprecipitation and western blotting

For immunoprecipitation experiments, cells were lysed as previously described. 300*μ*g protein (in 300 *μ*L total volume) per sample was pre-cleared with Protein A/G beads. The beads were pelleted and supernatant was incubated with Raf antibody (3*μ*L/sample) and beads overnight. All incubations included protease and phosphatase inhibitors in M-PER used for lysis with constant rotation at 4*°*C. Bead/antibody/protein slurries were then washed and subjected to standard SDS-PAGE analysis as previously described^17^. All antibodies were purchased from Cell Signaling (Boston, MA) with the exception of *α*-p621 Raf which was purchased from Biosource/Invitrogen (Carlsbad, CA), and *α*-CK2 from BD Biosciences (San Jose, CA).

### Morphology assessment

Untreated and ATRA-treated HL-60 cells were collected after 72 hr and cytocentrifuged for 3 min at 700 rpm onto glass slides. Slides were air-dried and stained with Wright’s stain. Slide images were captured at 40X (Leica DM LB 100T microscope, Leica Microsystems).

## Acknowledgements

We acknowledge the financial support to J.V. by the National Science Foundation CAREER (CBET-0846876) for the support of R.T. and H.J. In addition, we acknowledge support to A.Y. from the National Institutes of Health (CA 30555, CA152870) and a grant from New York State Stem Cell Science. The work described was also supported by the Center on the Physics of Cancer Metabolism through Award Number 1U54CA210184-01 from the National Cancer Institute. The content is solely the responsibility of the authors and does not necessarily represent the official views of the National Cancer Institute or the National Institutes of Health.

## Author contributions statement

J.V and A.Y directed the study. R.T, H.J, R.B and J.C conducted the cell culture measurements. J.V, R.B, W.D, K.R and A.S developed the reduced order HL-60 models and the parameter ensemble. W.D and J.V analyzed the model ensemble, and generated figures for the manuscript. The manuscript was prepared and edited for publication by W.D, R.B, A.Y and J.V. All authors reviewed this manuscript.

## Availability of the model code

The signaling and gene expression model equations, and the parameter estimation procedure, were implemented in the Julia programming language. The model equations were solved using the ODE23s routine of the ODE package64. The model code and parameter ensemble is available under an MIT software license and can be downloaded from http://www.varnerlab.org.

